# lcUMAPtSNE: Use of non-linear dimensionality reduction techniques with genotype likelihoods

**DOI:** 10.1101/2024.04.01.587545

**Authors:** Kerem Uzel, Christine Grossen, F. Gözde Çilingir

## Abstract

1. Understanding population structure is essential for conservation genetics, as it provides insights into population connectivity and supports the development of targeted strategies to preserve genetic diversity and adaptability.
2. T-distributed stochastic neighbor embedding (t-SNE) and uniform manifold approximation and projection (UMAP) have proven effective for revealing population genetic structures in human and model organisms using hard-called genotypes, but their application in wild species using genotype likelihoods from low coverage sequencing (as a cost-saving measure) remains underexplored.
3. Here, we present a Jupyter Notebook-based workflow that facilitates the use of UMAP and t-SNE on genotype likelihood-derived principal components.
4. This workflow is demonstrated using medium to low-coverage whole-genome sequencing data from scimitar-horned oryx, which has been reintroduced into the wild and faces multiple conservation challenges.
5. Detailed guidance on hyperparameter tuning and practical implementation is also provided, enhancing the application of these methods in wildlife genetics to potentially support biodiversity conservation.

## 1. Background

Population genetics seeks to understand genetic relationships within and between populations, informing strategies to preserve biodiversity and adaptability. Key applications include estimating population size, inferring demographic history, identifying population structure, and identifying loci that contribute to adaptive capacity or uncovering the genetic basis of reduced fitness in populations, all critical for effective conservation management, particularly for endangered species (Frankham, 2003; Hohenlohe et al., 2021).

The advent of next-generation sequencing has revolutionized population genetics by enabling genome-wide analyses, even for non-model species with low-coverage sequencing data. Low-coverage sequencing, combined with genotype-likelihood-based methods, provides a cost-effective approach to studying population structure and genetic diversity (Lou et al., 2021) while addressing uncertainties inherent in low-depth data (Korneliussen et al., 2014; Nielsen et al., 2011, 2012).

Dimensionality reduction techniques like principal component analysis (PCA) are widely used to visualize population structure. PCA simplifies genomic data by capturing major variance components, making it a foundational tool in genomics (Patterson et al., 2006). However, PCA’s linear assumptions may overlook non-linear genetic patterns (Alanis-Lobato et al., 2015), prompting the development of alternative techniques like t-distributed stochastic neighbor embedding (t-SNE) (van der Maaten & Hinton, 2008) and uniform manifold approximation and projection (UMAP) (McInnes et al. 2018). These non-linear methods can better capture complex relationships, especially when applied after PCA, to reduce computational demands and enhance interpretability (Kobak & Berens, 2019; Kobak & Linderman, 2021). Subsequently, top PCs are used as the input for further dimensionality reduction using t-SNE or UMAP (referred to as PCA-t-SNE and PCA-UMAP, respectively) (Diaz-Papkovich et al., 2019; van der Maaten & Hinton, 2008). In scenarios where genotype data is unavailable, such as in low-coverage whole-genome sequencing, PCs computed based on genotype likelihoods can be effectively incorporated into t-SNE and UMAP analyses. This adaptability further underscores the versatility of these dimensionality reduction techniques in various genomic data contexts.

Here, we introduce a Jupyter Notebook-based workflow, lcUMAPtSNE, that leverages genotype likelihood-based principal components for further dimensionality reduction using t-SNE and UMAP. This tool is demonstrated with medium to low-coverage whole-genome sequencing data from scimitar-horned oryx, a species with critical conservation needs. The workflow includes detailed guidance on parameter optimization and implementation, enabling ecologists and conservationists to apply these non-linear dimensionality reduction techniques in wildlife genetics effectively.

## 2. Description of the workflow

Our Jupyter Notebook (Kluyver et al., 2016) based workflow uses Python v3.11.4 (Van Rossum & Drake, 2009) and a set of Python libraries to perform t-SNE and UMAP analyses. The workflow begins by decomposing the covariance matrix output of PCAngsd, which incorporates genotype likelihoods and accounts for missing data while reflecting the genetic variance structure of the samples (Meisner & Albrechtsen, 2018). The covariance matrix is decomposed using NumPy (v1.24.4) (Harris et al., 2020) to extract eigenvectors and eigenvalues. Then, the principal components are computed by projecting the covariance matrix onto its eigenvectors, which are sorted in descending order of their eigenvalues. Since establishing a set of minimum and maximum number of available PCs was suggested (Diaz-Papkovich et al., 2019), our workflow incorporates a step for the guidance on the selection of principal components (PCs) for the input dataset that is performed by the ‘KneeLocator’ function from the ‘kneed’ Python package (v0.8.5) (Satopaa et al., 2011), which identifies the optimal number of PCs using a polynomial fit method to smooth the explained variance curve. This technique identifies the ‘elbow point’ by smoothing the PCs’ explained variance curve with polynomial interpolation and selecting the point where the rate of increase significantly diminishes (Fig. S1). Subsequently, user-defined principal components serve as input for dimensionality reduction via PCA-t-SNE and PCA-UMAP, utilizing the scikit-learn (v1.3.0) (Pedregosa et al., 2016) and umap-learn (v0.5.3) (McInnes et al. 2018) libraries, respectively. For clarity, we refer from here on to the techniques of PCA-t-SNE and PCA-UMAP as t-SNE and UMAP, respectively.

Hyperparameter optimization is a critical step in our workflow, conducted through a systematic grid search to fine-tune the settings for both t-SNE and UMAP. For t-SNE, we adjust the perplexity (*perp*), which influences the number of effective neighbors and typically ranges from 5 to 50 (van der Maaten & Hinton, 2008). For UMAP, we modify two parameters: the number of neighbors (*NN*, default 15) and the minimum distance (*MD*, default 0.1) (McInnes et al. 2018). Both *perp* and *NN* balance local versus global data structure representation, with smaller values focusing on local and larger ones on the global structure. The *MD* parameter controls data point separation, with lower values tightening clustering and higher values dispersing data points for global structure preservation.

Finally, all visuals of dimensionality reduction analyses are plotted with the seaborn library (v0.11.2) (Waskom, 2021). To ensure accessibility in visual data representation, we used a color palette generated by the online tool, which is available at https://medialab.github.io/iwanthue/.

To aid practical application, we provide several Jupyter Notebooks enabling users to input their covariance matrix, compute principal components, receive guidance on the minimum number of PCs to retain and apply t-SNE and UMAP techniques using various pre-determined hyperparameters to explore their effects on the dimensionality reduction outcomes. These notebooks were crafted to manage single or multiple input files (here genotype likelihood files) simultaneously, streamlining the analysis of diverse datasets with varied parameter configurations, thereby enhancing efficiency in handling extensive genomic datasets. Additionally, we provide a step-by-step guideline for users to run lcUMAPtSNE Jupyter Notebooks and explore their own datasets.

## 3. Example application: scimitar-horned oryx

The scimitar-horned oryx (*Oryx dammah*) is a large antelope that once roamed widely across North Africa (Bertram, 1988). However, the iconic animals experienced a precipitous decline in the 20th century due to drought, hunting, and land-use competition (Dixon et al., 1991), leading to their extinction in the wild (IUCN SSC Antelope Specialist Group, 2016). Before their disappearance, captive breeding began with fewer than 100 oryxes from Chad, expanding the ex-situ global population to about 15,000 individuals (Gilbert, 2019). While 1,000 are in coordinated breeding programs, many reside in places with minimal genetic management (Humble et al., 2023). Due to the success of reintroduction programs in Chad, the species was recently downlisted to Endangered by the IUCN Red List (IUCN SSC Antelope Specialist Group, 2023). To understand the genetic consequences of different conservation management strategies, (Humble et al., 2023) sequenced the whole genomes of 49 oryxes from populations with varying genetic management levels, notably from EAZA Ex Situ Programmes (EEP, n = 8), USA (n=17), and two unmanaged collections in the United Arab Emirates (EAD A, n = 9 & EAD B, n = 15).

### 3.1. Whole-genome resequencing data analyses

We obtained the raw whole-genome re-sequencing data of 46 genetically unrelated scimitar-horned oryx individuals from (Humble et al., 2023) (NCBI BioProject PRJEB37295), which were also utilized as the final dataset in the PCA analysis presented in the referenced study ([Humble et al., 2023]; Table S1). Subsequent data processing was performed using the ATLAS Pipeline v7 (Link et al., 2017; Marchi et al., 2022; https://atlaswiki.netlify.app/atlas-pipeline), involving the Gaia, Rhea, and Perses workflows. In Gaia, individual sequencing data underwent quality trimming using TrimGalore v0.6.6 (https://www.bioinformatics.babraham.ac.uk/projects/trim_galore/) with default settings and was aligned to the *Oryx dammah* reference genome, SCBI_Odam_1.1 (NCBI RefSeq GCF_014754425.2) (Humble et al., 2020) employing BWA-MEM v0.7.17 (Li & Durbin, 2009) in paired-end mode. Within the Rhea workflow, local re-alignment was conducted using GATK v3.8 (McKenna et al., 2010) (Table S1). Subsequently, the Perses workflow facilitated the merging of forward and reverse reads into single reads per fragment using the tool ATLAS v0.9 (Link et al., 2017), avoiding pseudo-duplicated bases where original reads overlapped.

After quality filtering, we retained an average of 99% of the raw sequencing data coming from 46 scimitar-horned oryx individuals. After mapping and deduplication, an average of 81% (range: 77-87%) of these high-quality reads mapped to the reference genome. The approximate individual-level coverage varied from ∼4.6x to ∼20.6x (Table S1).

We implemented downsampling with ATLAS (Link et al., 2017) (task=downsample) to standardize the coverage across individuals, setting high-coverage samples to an average coverage of 6x while preserving the original coverage levels for lower-coverage individuals in accordance with the methodology outlined in the cited study. We designated the primary dataset as “SO_6x”. In this set, we maintained the 20 samples with coverage below the 6x average and downsampled the remaining 26 to the target of ∼6x (Table S1). For subsequent analyses to explore the effect of lower sequencing coverage, we created “SO_2x” and “SO_0.5x”, where we downsampled all samples to about 2x and 0.5x, respectively.

### 3.2 Genotype likelihood estimation and PCA

We estimated genotype likelihoods using ANGSD v0.940 (Korneliussen et al., 2014) based on the three distinct datasets described above, each characterized by varying levels of average coverage (6x, 2x, 0.5x). Following the methodology of (Humble et al., 2023), we employed ANGSD (Korneliussen et al., 2014) using the GATK model (-GL 2) to infer major and minor alleles (-doMajorMinor 1, -doMaf 1). We restricted this analysis to the 28 chromosome-length autosomes (Table S1) and included only regions with Phred quality and mapping scores exceeding 30. We utilized properly paired (-only_proper_pairs 1) and unique reads (-uniqueOnly 1) while retaining only biallelic sites (-skipTrialleleic 1). Sites with read coverage in less than 60% of the samples were excluded (-minInd 30, allowed missingness 60%), and we retained only polymorphic sites with a genotype likelihood p-value less than 1e-6 (-SNP_pval 1e-6). We applied thinning for the variant sites, ensuring a minimum distance of 1Kb between two sites with a custom bash script. Lastly, we performed the principal component analysis with PCAngsd v1.11 (Meisner & Albrechtsen, 2018) by using default settings where the minor allele frequency cut-off is 5%.

SO_6x (dataset with coverage 4.6x-6x) yielded a total of 1,415,641 variant sites with a minor allele frequency (MAF) greater than 0.05 and a minimal pairwise distance of 1kb. SO_2x yielded 1,516,230, and SO_0.5x yielded 23,129 variant sites with MAF > 0.05 and distance > 1kb. The PCA performed was insensitive to coverage, with all three coverage datasets mirroring the PCA results reported in Humble et al. (2023), where PC1 separated the genetically unmanaged population EAD_B from the rest, PC2 separated the other genetically unmanaged population EAD_A from the cluster of genetically managed populations (EEP + USA) and PC3 separated the genetically managed EEP population from the others (Figs 1A-C, Figs S2A-C). The cumulative variance explained by PC1 and PC2 for the SO_6x, SO_2x, and SO_0.5x datasets were 21.9%, 20.3%, and 19.2%, respectively.

**Fig. 1.**
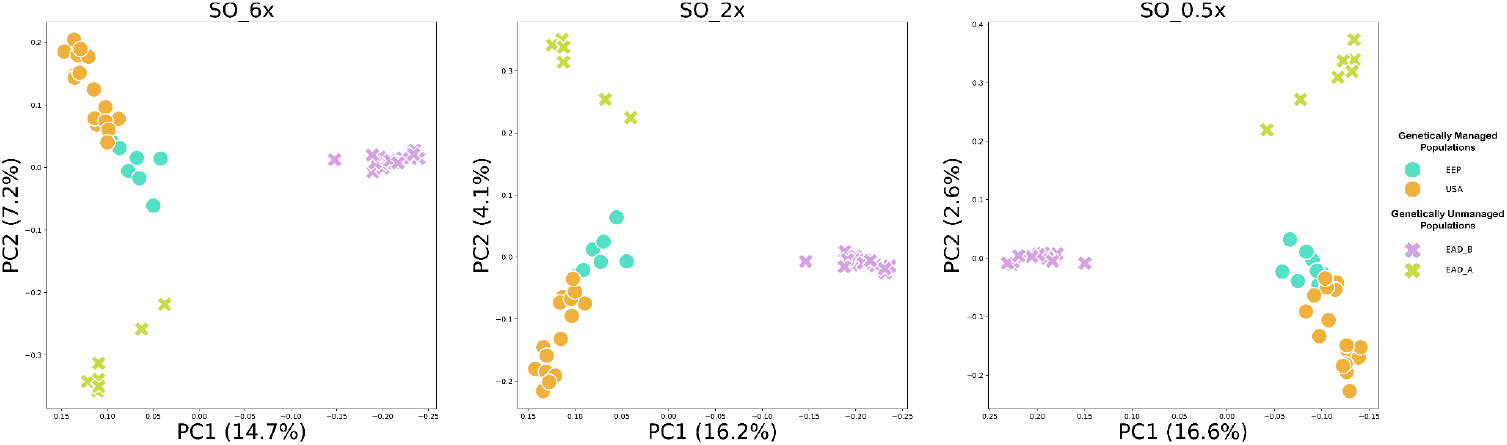
The PCA plot of the SO_6x (A), SO_2x (B), and SO_0.5x (C) datasets using the first two PCs. Each dot represents a single individual.

### 3.3 t-SNE and UMAP applications using lcUMAPtSNE workflow

We tested various combinations of *perp, NN*, and *MD* values across three datasets. For t-SNE, we used *perp* values of 5, 10, and 23 (up to half the total number of individuals), while for UMAP, we employed *NN* values of 5, 10, and 23, along with *MD* values of 0.01, 0.1 (the default), and 0.5. These parameters were combined with a set of top PCs where the elbow point approach suggested using 5 to 46 PCs for the three datasets. For instance, in the SO_6x dataset, we applied 6 (the elbow point) and 46 PCs (the maximum available, as shown in Fig. S1A); similarly, for both SO_2x and SO_0.5x datasets, we used 5 and 46 PCs (Figs S1B & C).

The minimum (N=5) and maximum number of PCs (N=46) incorporated in the t-SNE and UMAP analyses of SO_6x, SO_2x, and SO_0.5x corresponded to 35.3 and 99.8%, 26.6 and 99.3%, 24.0 and 98.8% cumulative variance explained, respectively (Fig. S1). To ensure clarity, we present in the main figures results acquired using the mid-range parameters selected for t-SNE and UMAP (*perp* 10, *NN* 10, *MD* 0.1) coupled with the top PCs obtained with the elbow method unless stated otherwise, as these parameters effectively capture the overall trends observed across the hyperparameter space we explored. See the Supplementary Material for a comprehensive overview of the parameter space we explored.

The t-SNE and UMAP analyses of all three datasets of varying coverages, using the mid-range parameters) confirmed four groups representing the four sampled populations (Fig. 2 & Figs 3A, B; Figs S3-8). All three datasets successfully differentiated the two genetically managed groups, whereas the top two principal component projections (PC1 vs PC2) from the PCA did not achieve this distinction (Fig. 2; Figs S3-6). The combination of PC1 and PC3 did, but then the resolution of EAD_A from EEP was lost (Fig. S2). For both datasets, as anticipated, the cohesiveness of the clusters improved with combinations of local parameters. Lowering the *perp, NN*, or *MD* values led to clustering at a finer scale, while increasing these values helped to visualize more global patterns, suggesting higher genetic similarity between (and leading to clustering of) one unmanaged population (EAD_B) and the two managed populations than each of these with the other genetically unmanaged population (EAD_A) (Figs S3–6). At 6x and 2x, the increase in the number of PCs did not influence the clustering patterns obtained using either technique (Figs S4 & S6).

**Fig. 2.**
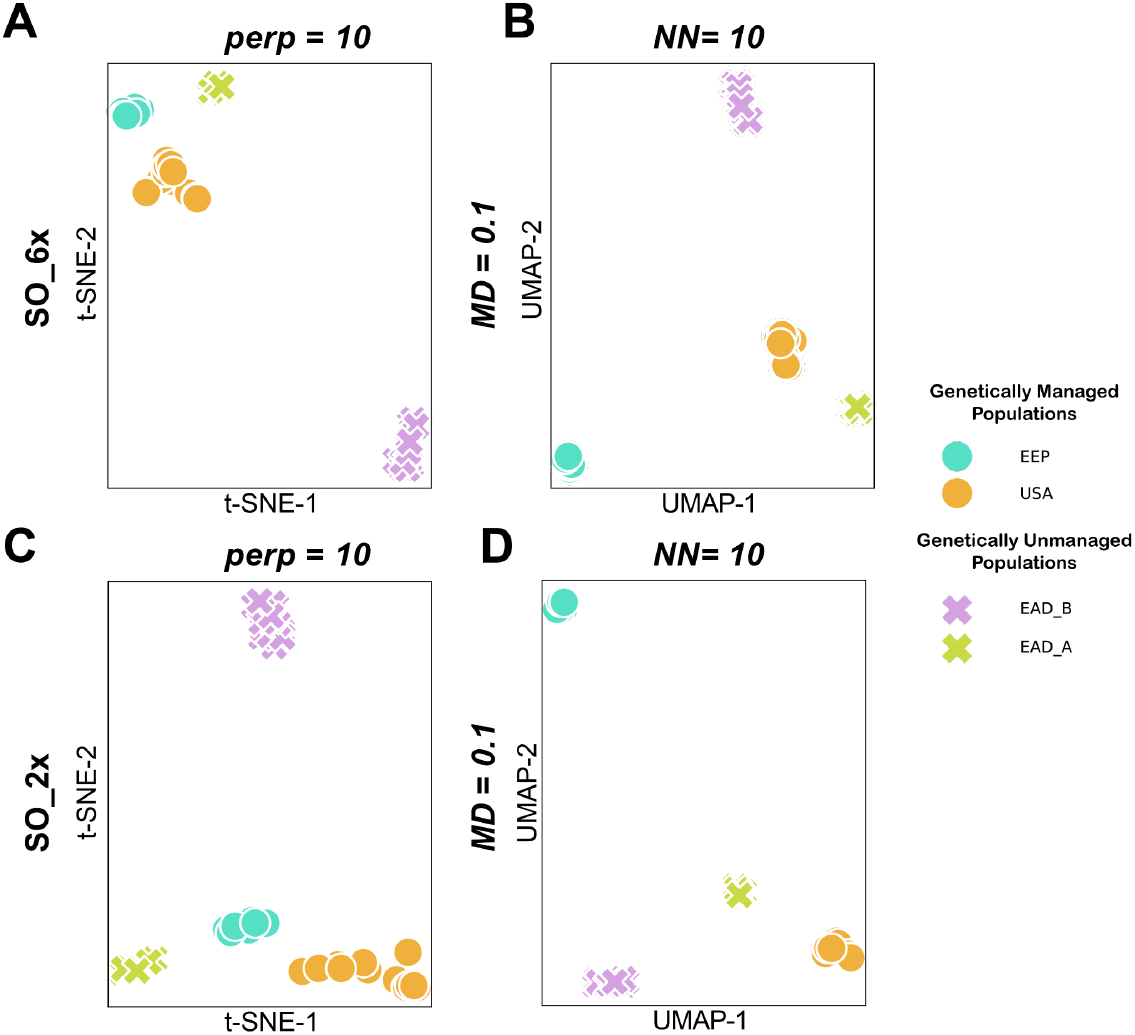
t-SNE and UMAP representations of SO_6x (A, B) with 6 PCs and of SO_2x (C, D) with 5 PCs (minimum number of significant PCs based on elbow method). Each dot represents a single individual.

**Fig. 3.**
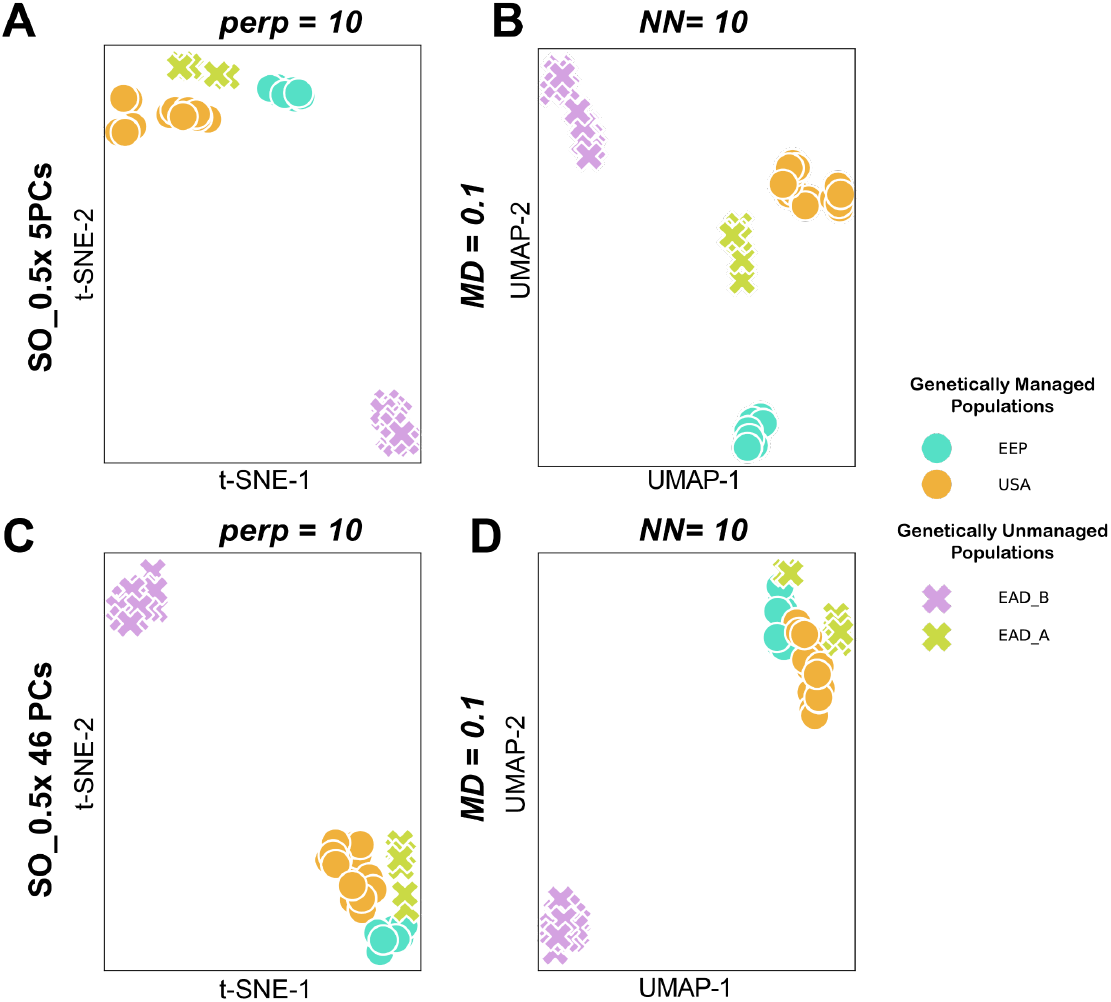
t-SNE and UMAP representations of SO_0.5x with 5 PCs (minimum number of significant PCs, A, B) and with 46 PCs (maximum number of PCs, C. D). Each dot represents a single individual.

For SO_0.5x, both t-SNE and UMAP also separated the two genetically managed populations (Figs 3A & B; Figs S7 & S8). However, t-SNE analyses provided poorer local structure when compared to UMAP (Figs 3A & C). Notably, when we increased the number of PCs, the clustering efficacy of UMAP was reduced as opposed to the trends observed with SO_6x and SO_2x (Figs 3C & D; Fig. S8), likely due to the added uncertainty of genotype likelihoods with 0.5x (Meisner & Albrechtsen, 2018).

## 4. Discussion

UMAP and t-SNE have been effectively applied to analyze population genetic structure across various species, including plants, invertebrates, and humans (*Anopheles gambiae* 1000 Genomes Consortium, 2020; Černý et al., 2023; Chyleński et al., 2019; Diaz-Papkovich et al., 2019; Fu et al., 2022; Halldorsson et al., 2022; Ma et al., 2021; Margaryan et al., 2020; Schmidt et al., 2020, 2021; Sengupta et al., 2021; Simon et al., 2021; Sohail et al., 2023). Despite their proven utility, these non-linear dimensionality reduction methods are underutilized in conservation genomics, particularly with genotype likelihoods from low-coverage sequencing—a gap that presents a significant opportunity for advancing research in this field.

We introduce a Jupyter Notebook-based workflow, lcUMAPtSNE, which allows users to apply PCA-initialized UMAP and t-SNE to genotype likelihoods. Demonstrated with the Scimitar-horned oryx dataset across three coverage levels (6x, 2x, 0.5x), this workflow effectively discerns populations and provides enhanced resolution beyond PC3, summarized in a single visualization.

While default parameters like perplexity (perp), number of neighbors (NN), and minimum distance (MD) work well in large human datasets (Diaz-Papkovich et al., 2019, 2020), smaller sample sizes in conservation genomics require a customized approach. Our lcUMAPtSNE workflow facilitates this customization through a grid search, allowing users to explore parameter settings systematically. Importantly, using the elbow method to determine the optimal number of PCs avoids incorporating excess noise, optimizing the analysis for different data characteristics. With the provided online step-by-step guidelines, users can explore their own datasets.

lcUMAPtSNE is an easy-to-implement tool. Therefore, users can supplement their PCAs with non-linear dimensionality reduction techniques even for lower coverage datasets by utilizing genotype likelihoods. It is crucial to recognize that both linear and non-linear approaches to dimensionality reduction can result in data distortion, each presenting unique benefits and challenges. As such, careful handling of data and biologically relevant interpretation of the results are imperative.

## Acknowledgments

We would like to thank all WSL Ecological Genetics Group members who provided their comments on the manuscript. We extend our gratitude to the authors who have made their genomic datasets available for open access. Specifically, we thank Emily Humble et al. for the scimitar-horned oryx dataset and all others who contributed to the resources that significantly supported this research. F. Gözde Çilingir was funded by the University of Zurich Postdoc Grant #FK-22-109.

## Data accessibility statement

All raw sequencing data used in this study were downloaded from public databases, and no new data were generated. All intermediate files we produced using these datasets, all bioinformatic codes used for generating the results we presented, and guidelines are available at https://github.com/fgcilingir/lcUMAPtSNE.

## Conflict of interest

The authors have no conflict of interest to declare.

## Author contributions

F. Gözde Çilingir conceived the project idea and designed the study with input from Christine Grossen and Kerem Uzel. F. Gözde Çilingir and Kerem Uzel performed the bioinformatic analyses and prepared the online guidelines. Kerem Uzel and F. Gözde Çilingir wrote the manuscript with substantial input from Christine Grossen. All authors reviewed the manuscript.

## Notes

https://github.com/fgcilingir/lcUMAPtSNE

### Competing Interest Statement

The authors have declared no competing interest.

### Summary of Updates

In this revised manuscript, we have updated the author contribution order, title, and scope to emphasize the application of our Jupyter Notebook-based workflow. This version focuses on demonstrating the workflow using the Scimitar-horned oryx dataset only. We have also revised the introduction and the discussion/concluding remarks to better align with the new focus.

