## Supplementary figures and images for "lcUMAPtSNE: Use of non-linear dimensionality reduction techniques with genotype likelihoods"

### Fig. S1

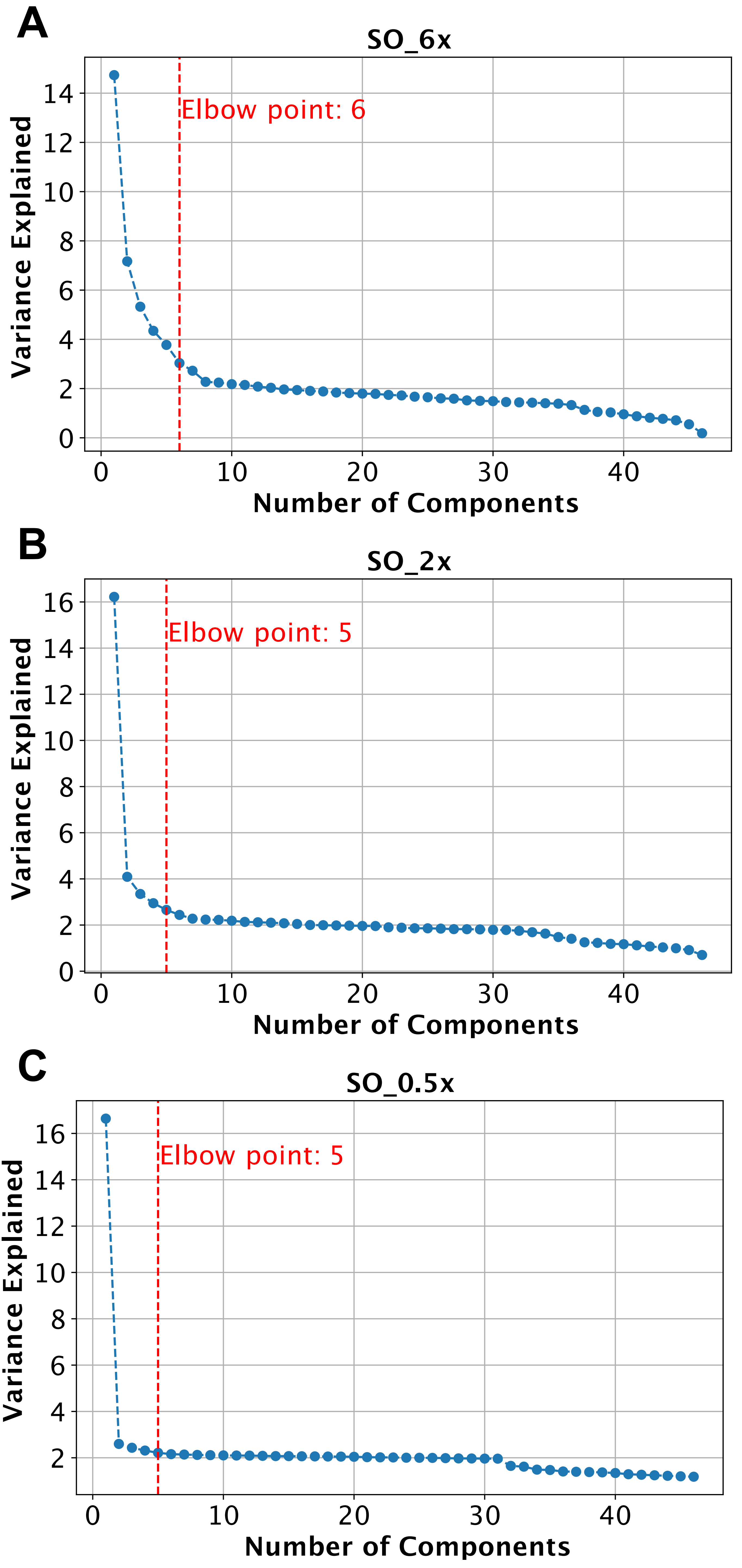

### Fig. S2

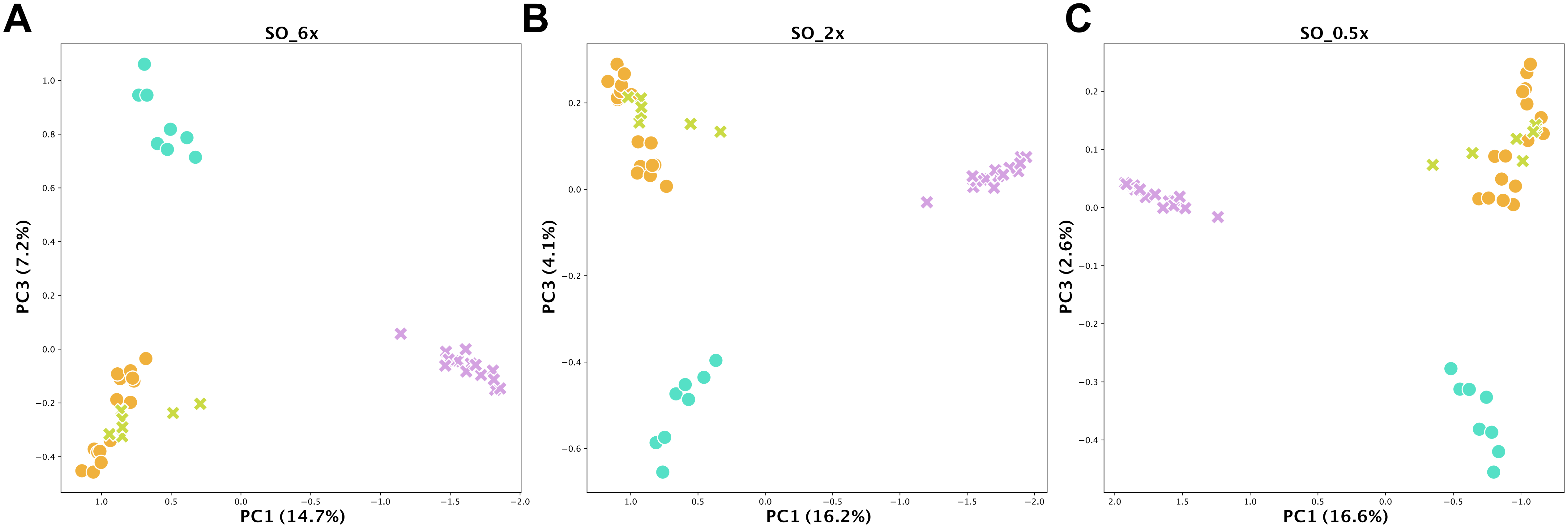

### Fig. S3

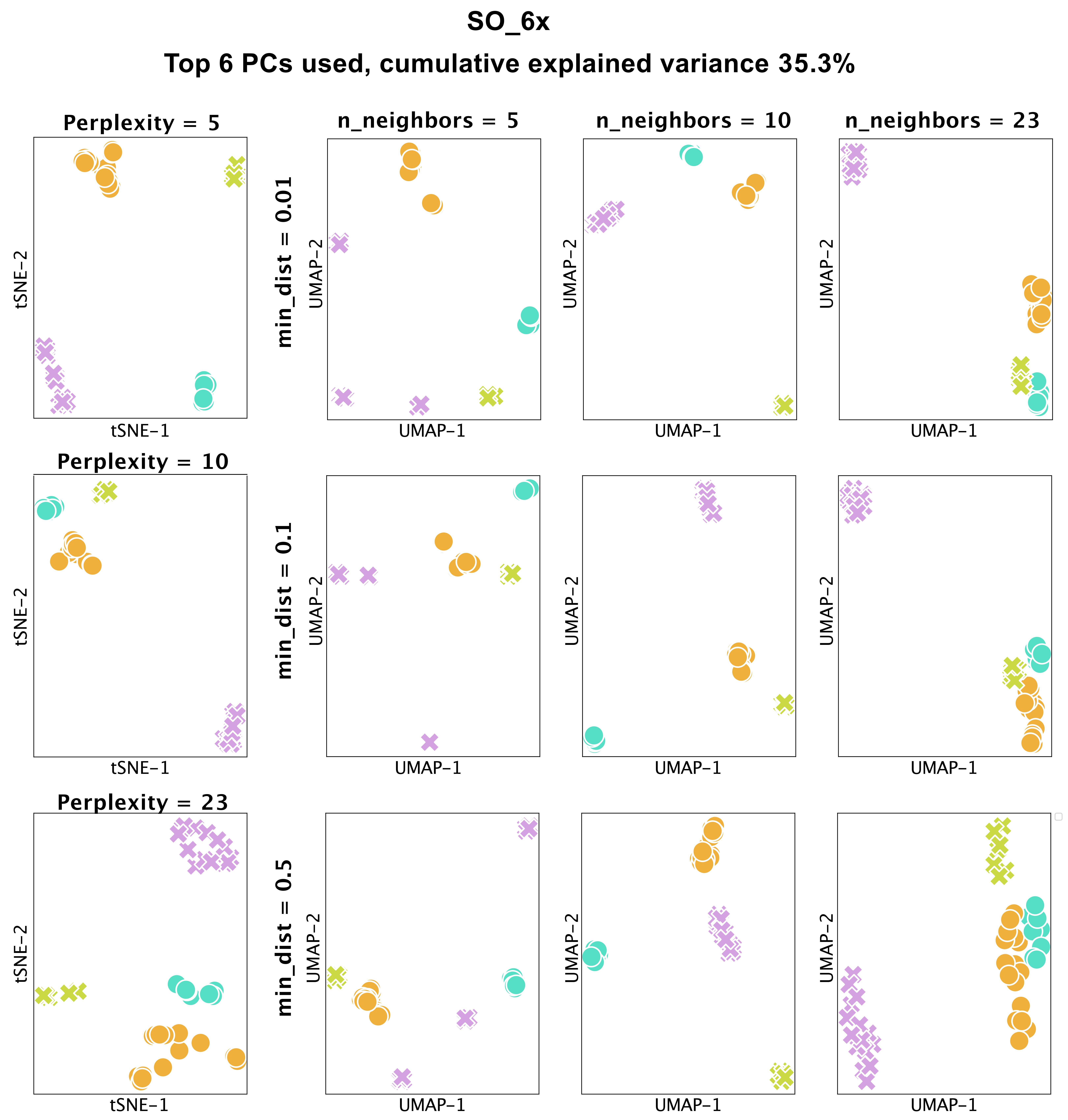

### Fig. S4

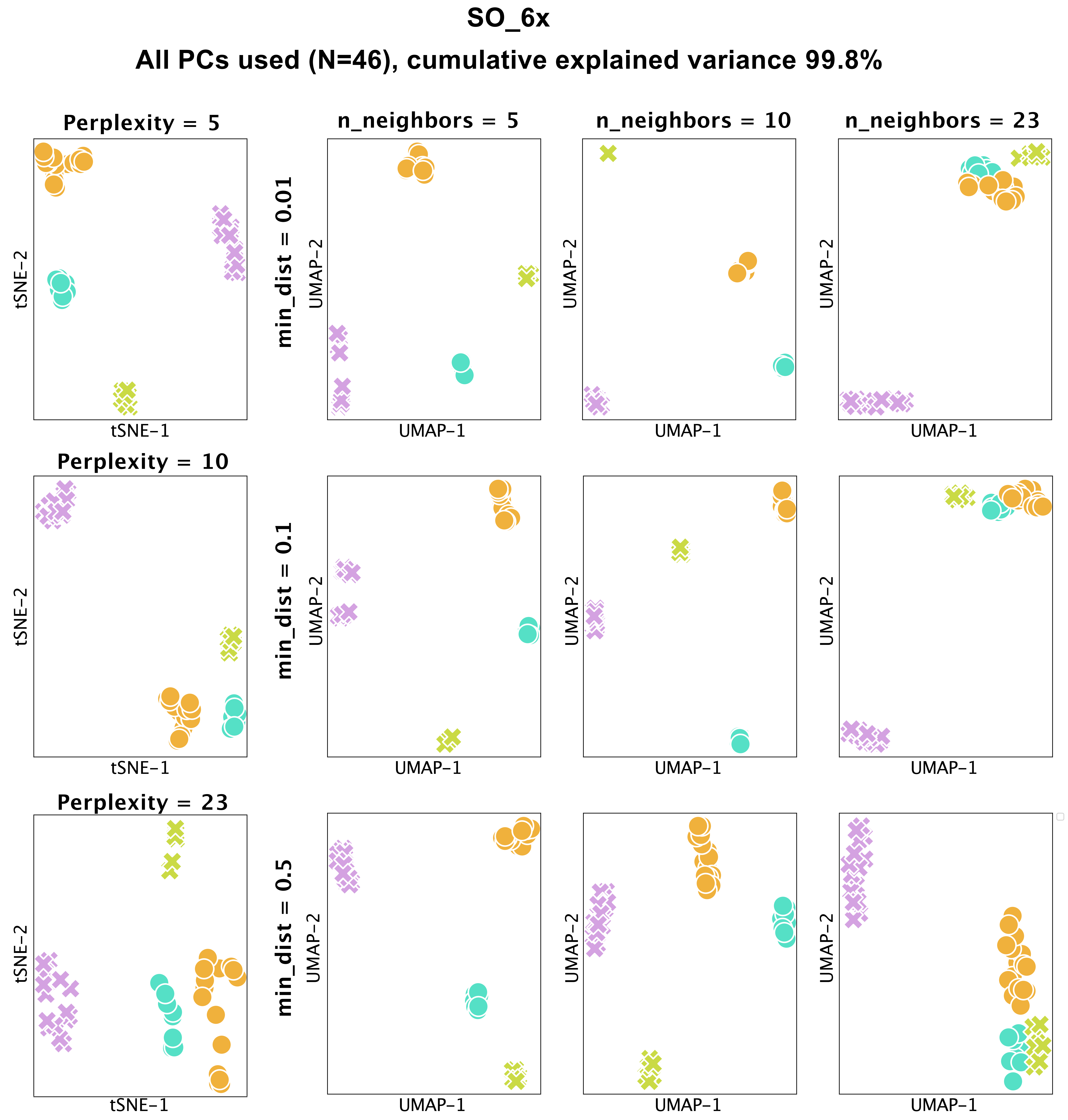

### Fig. S5

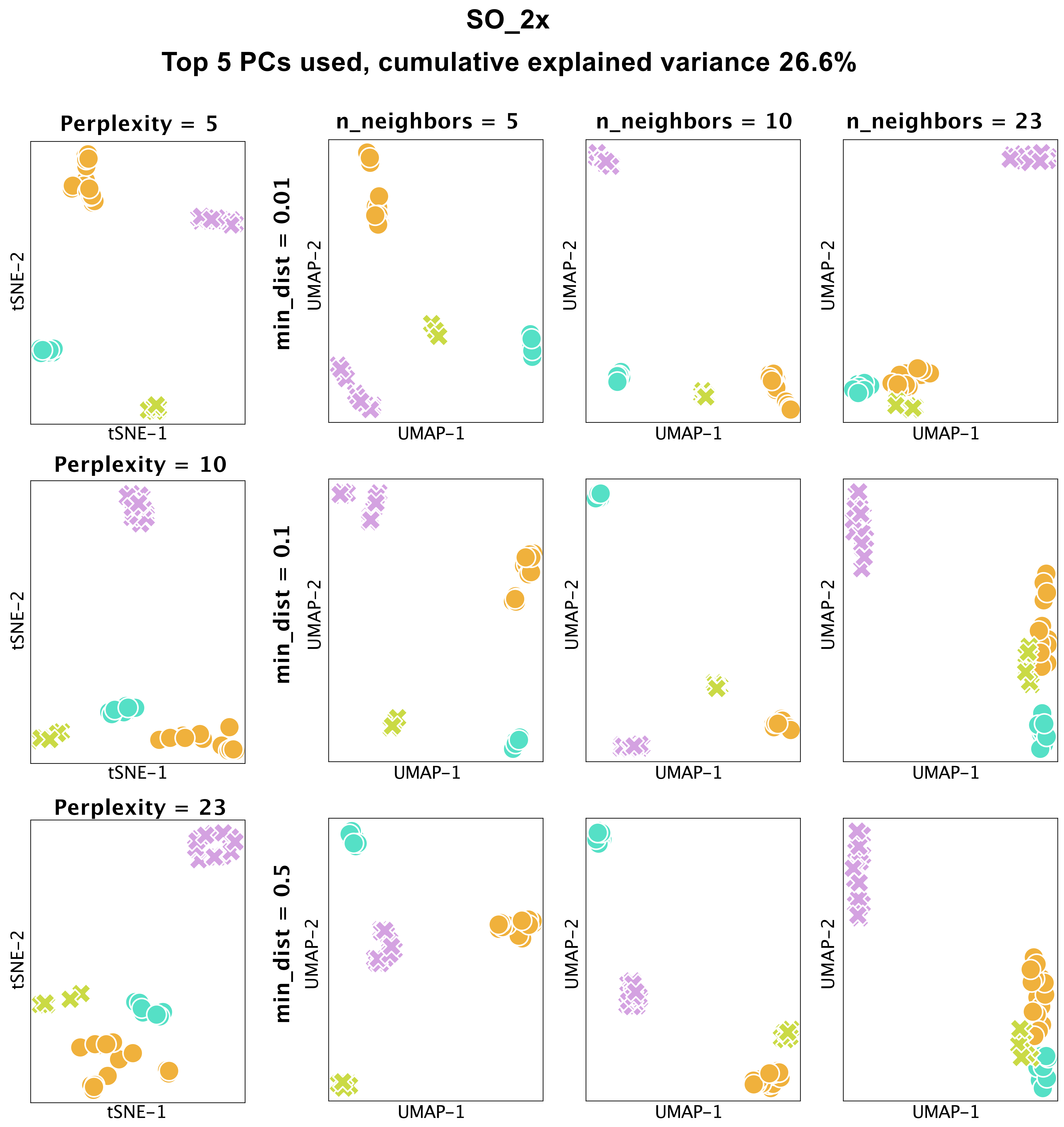

### Fig. S6

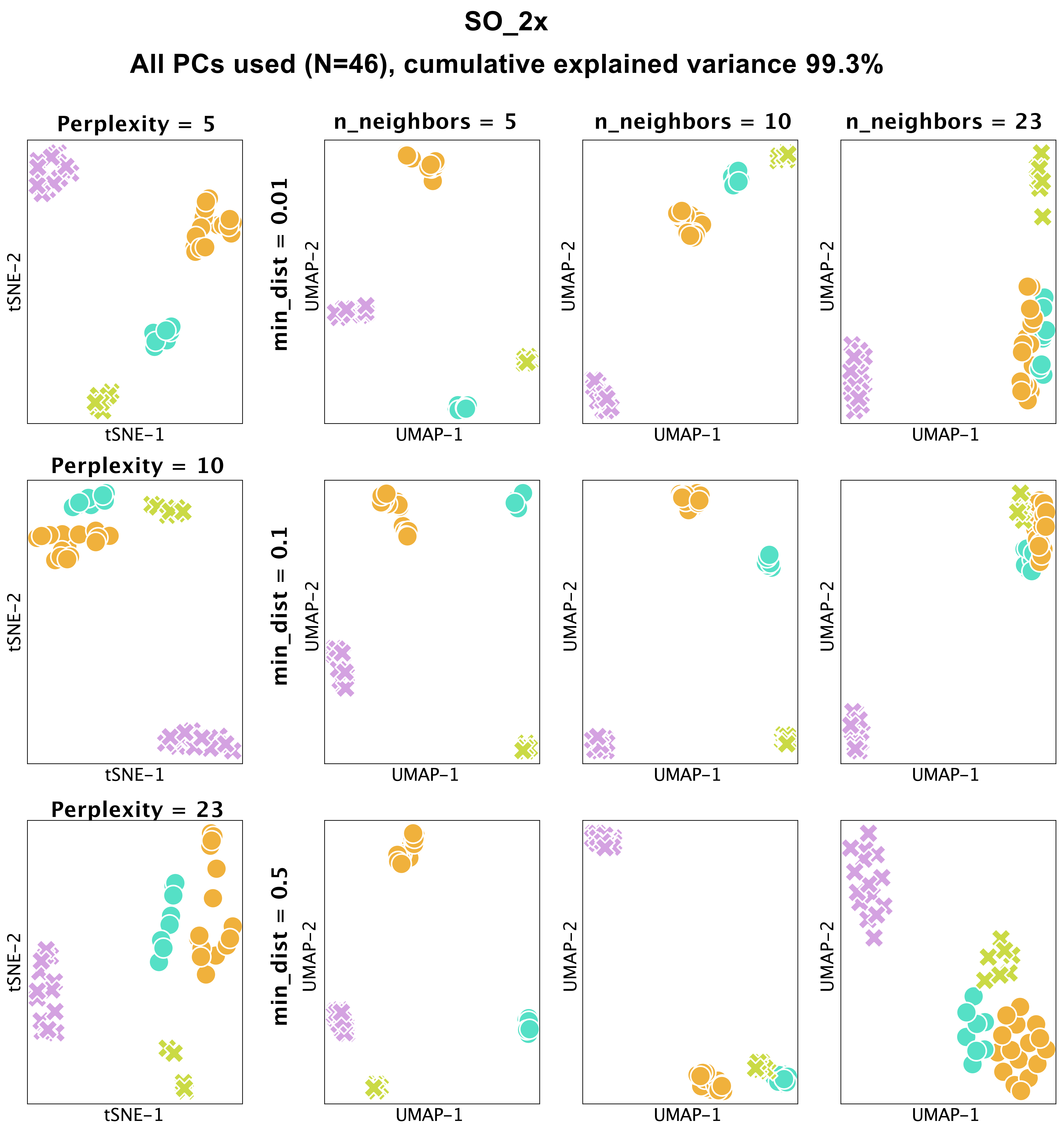

### Fig. S7

**SO_0.5x**

**Top 5 PCs used, cumulative explained variance 24.0%**

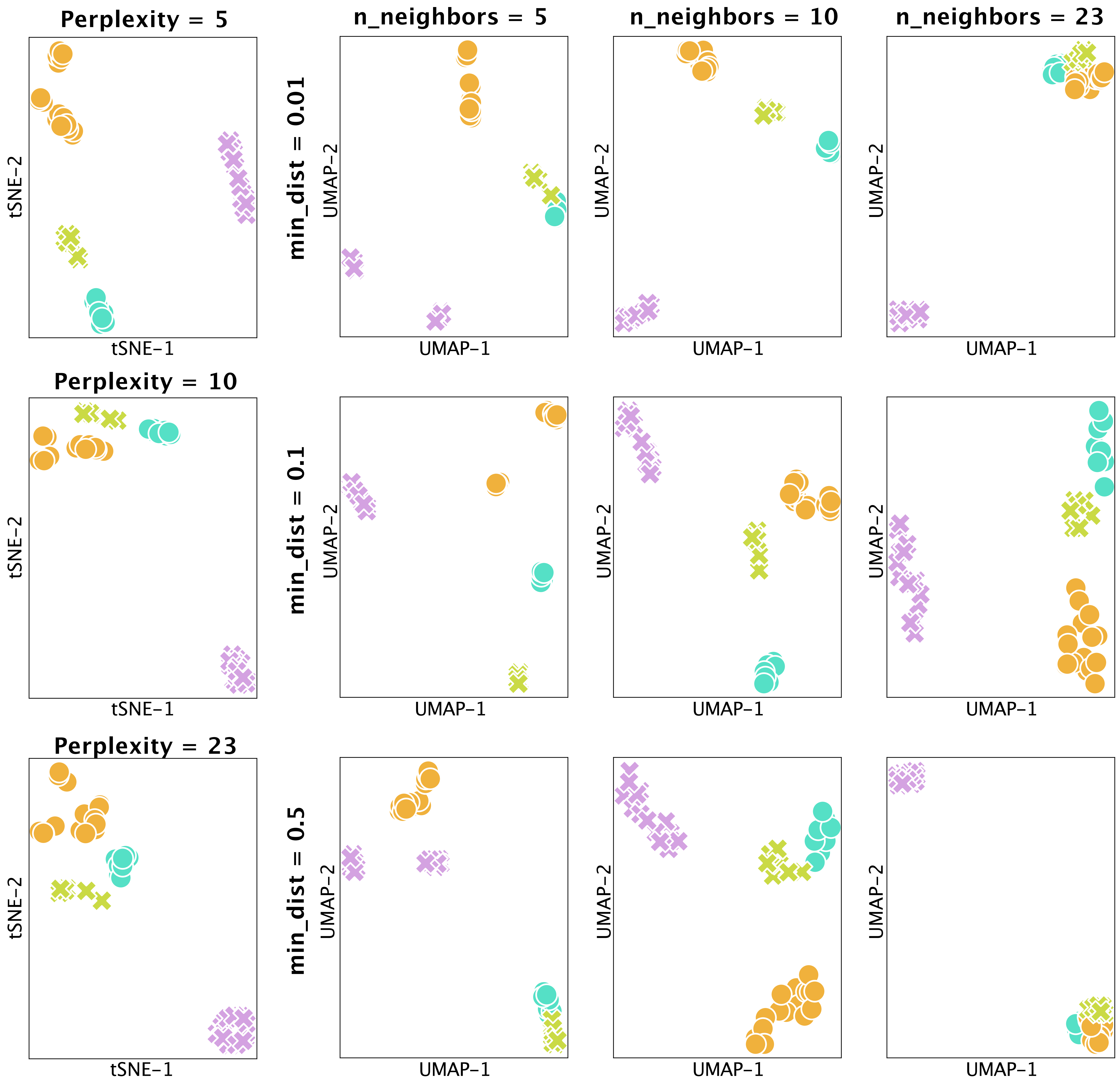

### Fig. S8

# SO_0.5x

#### All PCs used (N=46), cumulative explained variance 98.8%

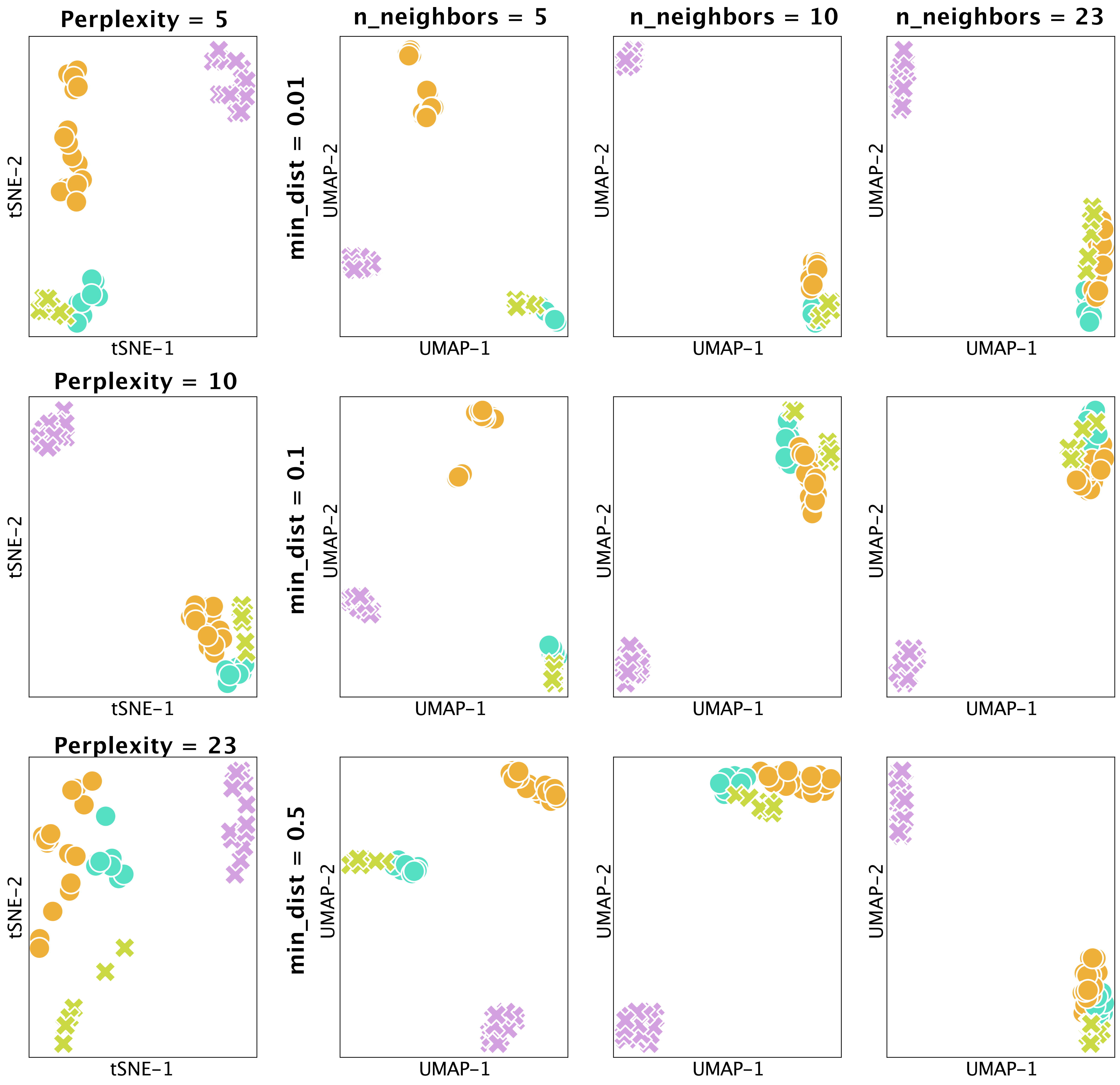
